# Ovulation sources coagulation protease cascade and hepatocyte growth factor to support physiological growth and malignant transformation

**DOI:** 10.1101/2021.06.16.448625

**Authors:** Hsuan-Shun Huang, Pao-Chu Chen, Sung-Chao Chu, Ming-Hsun Lee, Chi-Ya Huang, Tang-Yuan Chu

**Author notes:** First authors with equal contribution (Hsuan-Shun Huang, Pao-Chu Chen). **To whom correspondence should be addressed:**, Tang-Yuan Chu, MD, PhD, Department of Obstetrics and Gynecology, Buddhist Tzu Chi General Hospital, 707, Section 3, Chung-Yang Road, Hualien, Taiwan, ROC.

## Abstract

The fallopian tube fimbrial epithelium (FTE), which is exposed to the follicular fluid (FF) contents of ovulation, is regarded as the main origin of ovarian high-grade serous carcinoma (HGSC). Previously, we found that growth factors in FF, such as IGF2, are responsible for the malignant transformation of FTE. However, ovulation is a monthly transient event, whereas carcinogenesis requires continuous, long-term exposure. Here, we found the transformation activity of FF sustained for more than 30 days after drainage into the peritoneal fluid (PF). Hepatocyte growth factor (HGF), activated through the ovulation injury-tissue factor–thrombin–HGFA–HGF cleavage cascade confers a sustained transformation activity to FTE, HGSC. Physiologically, the high reserve of the coagulation-HGF cascade sources a sustained level of HGF in PF, then to the blood circulation. This HGF axis promotes the growth of the corpus luteum and repair of tissue injury after ovulation.

**Graphical Abstract:** 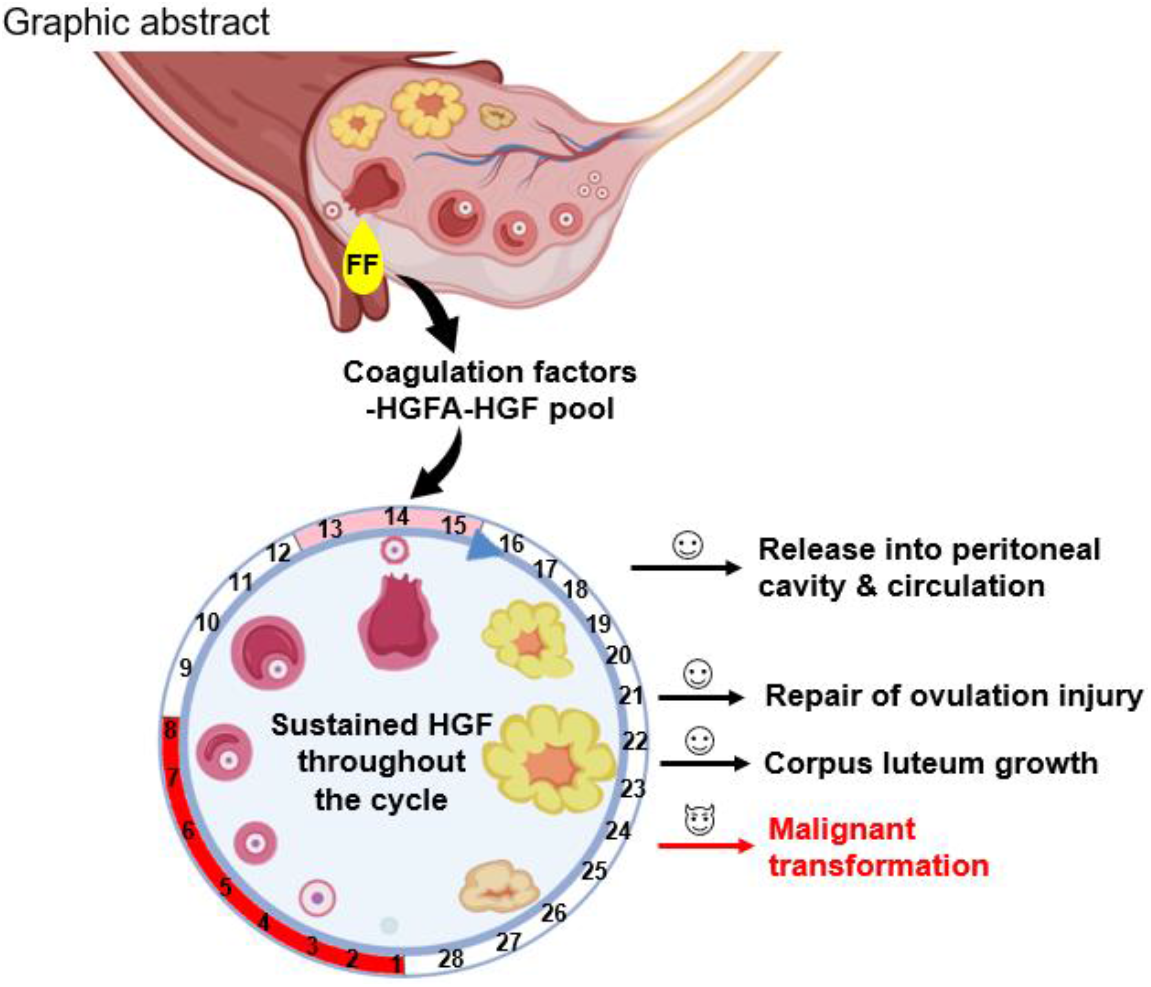

The fluid content of the ovulating ovarian follicle (follicular fluid [FF]) exerts a long-lasting transformation effect that is sustained after mixing with peritoneal fluid (PF). The coagulation protease cascade and hepatocyte growth factor (HGF) are responsible for the sustained activity. The high abundance of cascade proteins and pro-form HGF activator and HGF in FF/PF supports the constant supply of active-form HGF lasting for a whole month, covering the entire ovulation cycle. This sustained HGF activity promotes the proliferative repair of the ovarian surface and fallopian tube fimbrial epithelia as well as the growth of the corpus luteum. The same activity also inadvertently promotes the malignant transformation of the fimbrial epithelium, high-grade serous carcinoma, and epithelial ovarian cancer. In addition to the peritoneal cavity, ovulation sources HGF to blood circulation, which may serve an endocrine role.

**HIGHLIGHT:**

1. Ovulatory follicular fluid (FF) exerts a long-lasting transformation activity covering throughout the ovulation cycle.
2. The ovulation injury-coagulation proteases-hepatocyte growth factor (HGF) cascade is responsible for the sustained activity.
3. Ovulation sources HGF into the peritoneal cavity, then into the blood circulation.
4. This coagulation-HGF cascade promotes the transformation of fallopian tube epithelial cells and ovarian cancer cells.
5. Physiologically, it promotes the growth of the corpus luteum and injured epithelium after ovulation.

## INTRODUCTION

Every year, approximately 190,000 incident cases of ovarian cancer are diagnosed globally, resulting in 114,000 deaths (Bharwani et al., 2011). Ovarian cancer is the seventh most common cancer among women worldwide [https://gco.iarc.fr/today/home]. Epithelial ovarian cancer (EOC) accounts for 90% of ovarian cancer cases. In Western countries, more than 70% of EOC cases are high-grade serous carcinoma (HGSC)(Seidman et al., 2004). The majority of HGSC cases are diagnosed at stage III (~50%) or stage IV (~30%), with 5-year survival rates of 42% and 26%, respectively(Torre et al., 2018). Because of highly rapid intraperitoneal spread, early-stage HGSCs are both uncommon and difficult to diagnose. Common physiological changes in the morphology, size, and content of the ovary can obscure the diagnosis through imaging. Moreover, deep pelvic localization prohibits an easy histological diagnosis through tissue biopsy.

Recent studies have provided insights into the etiology and tissue of origin of HGSCs(Mei et al., 2021; Wu et al., 2020). The majority of HGSCs are derived from the secretory cells of the fallopian tube epithelium (FTE), especially at the fimbriae end (Levanon et al., 2008), which approaches the ovulating follicle to catch the oocyte while exposing itself to follicular fluid (FF) after ovulation.

Large-scale epidemiological studies and meta-analyses have indicated that the lifetime number of ovulation cycles is positively associated with ovarian cancer risk(Trabert et al., 2020; Wu et al., 1988). The use of oral contraceptives (OCs) to inhibit ovulation prevents the incidence of ovarian cancer in an early-onset, long-lasting, and dose-dependent manner(Greer et al., 2005; Havrilesky et al., 2013). In addition, profound protection against ovarian cancer was found in the physiological states of anovulation such as term pregnancy(Yang et al., 2007; Yen et al., 2003) and lactation (La Vecchia, 2017; Yen *et al.*, 2003). These findings suggest a carcinogenic effect of ovulation and the presence of carcinogens in ovulatory FF.

Previous studies from us have reported a potent transforming activity of FF. When repeatedly injected into the mammary fat pad of Trp53^−/−^ mice, FF induced early-onset lymphomas in injection sites in more than half of the mice(Hsu et al., 2019; Huang et al., 2015). In immortalized FTE cells (FTECs) with different alterations in *TP53* and *RB1* or *CCNE1*, FF induced the malignant phenotypes of anchorage-independent growth (AIG) and xenograft tumorigenesis(Hsu et al., 2021; Hsu *et al.*, 2019; Huang *et al.*, 2015). In addition, HGSC cells with either competent or defective homologous recombination repair are all vulnerable to the transforming effect of FF(Hsu *et al.*, 2021). A wide spectrum of malignant phenotypes is augmented by FF. The order of magnitude of these phenotypes is as follows: cell migration, AIG, invasion, peritoneum attachment, anoikis resistance, and proliferation(Hsu *et al.*, 2021).

Transforming agents present in FF remain unclear. Previously, we found IGF-axis proteins in FF, including IGF2, IGFBP2/6, and the lytic enzyme PAPP-A are largely responsible for the AIG and tumorigenesis activity of FF. Through IGF-1R/AKT/NANOG and IGF-1R/AKT/mTOR pathways, the FF–IGF axis promotes stemness activation, clonal expansion, and FTEC transformation(Hsu *et al.*, 2019; Wu *et al.*, 2020). However, the FF–IGF axis does not confer full transformation activity and phenotypes. Other transforming factors in FF remain to be identified. Moreover, the exposure of FF to the fimbria is only transient before it is diluted in the peritoneal fluid (PF), and this ovulatory exposure occurs only once monthly. The long-term transformation activity of FF after diluting into PF (named FF/PF) remains unknown.

In this study, we observed a long-lasting transformation effect of FF/PF throughout the ovulation cycle. Hepatocyte growth factor (HGF) and its receptor c-MET are responsible for the transformation effect. The reserve of the blood coagulation–HGFA–HGF axis proteins in FF is responsible for the sustained activity. Physiologically, this durable HGF activity may be responsible for the repair of postovulatory tissue damage and the growth of the corpus luteum.

## RESULTS

### Human PF exhibits prominent transformation activity after ovulation, which is sustained throughout the menstrual cycle

After ovulation, the contents of the ovarian follicle drain into the peritoneal cavity and mix into PF. We collected PF from 15 women with regular menstruation during laparoscopic surgery (Table 1). The time to ovulation was determined on the basis of the menstrual history and the date of the most recent and previous menstrual period. Ten and five PF samples were collected in the luteal and proliferative phases of the menstrual cycle, respectively. The AIG activity of each PF sample was determined by treating two immortalized fimbrial epithelial cell lines (FE25 and FT282-CCNE1) grown in soft agar. As shown in Figure 1, PF samples collected in the luteal phase had higher AIG activity than did those collected in the proliferative phase. However, no inhibitory effect was found using IGF1R inhibitors. A slow and steady decline in AIG activity was determined after ovulation (R^2^ = 0.3351; Fig. 1B); however, a considerable high AIG activity was maintained throughout the ovulation cycle.

**Fig 1.**
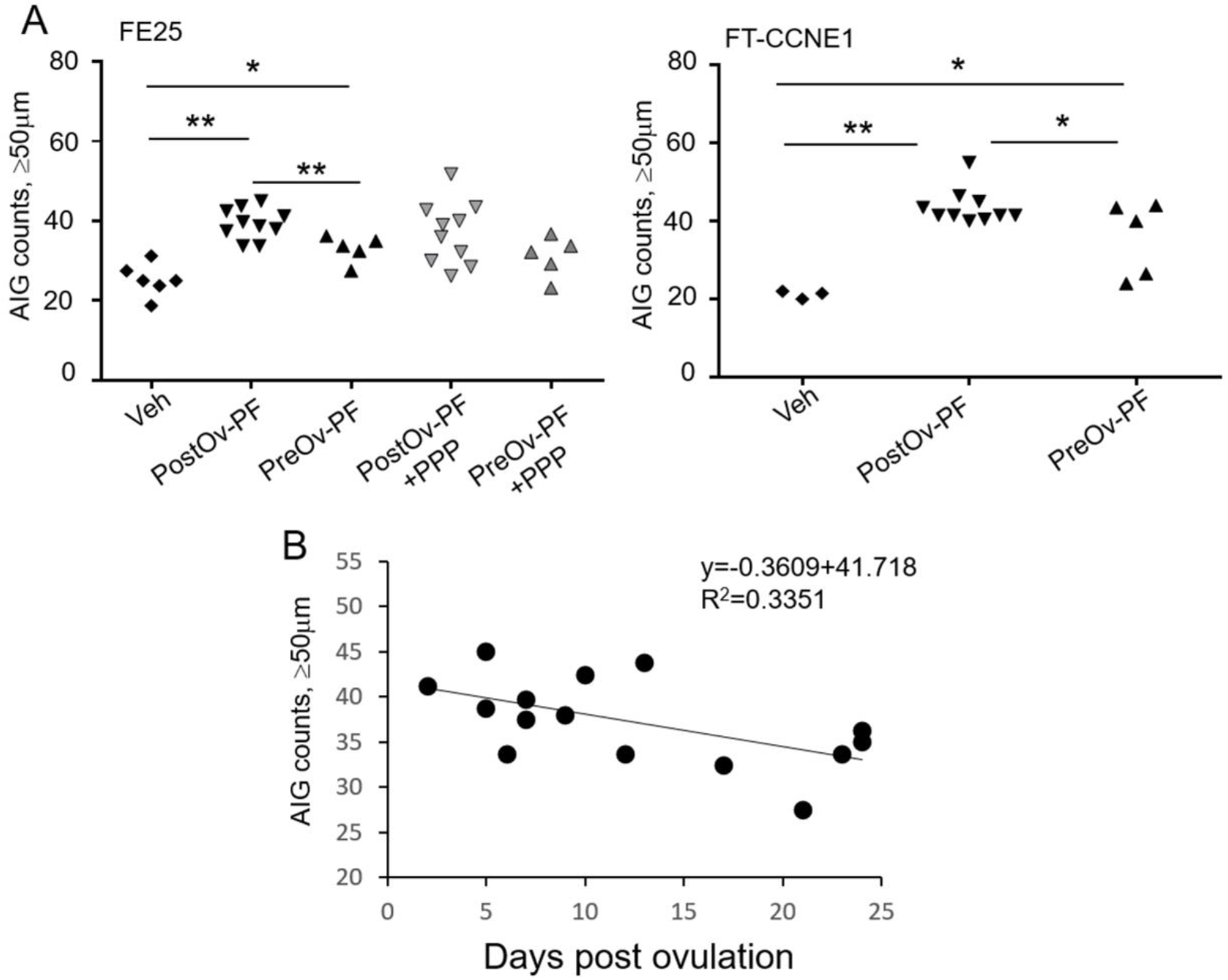
Sustained transformation activity of human peritoneal fluid with a slow decline after ovulation. (A) AIG colony counts of two FTE cell lines (FE25 and FT-CCNE) after treatment with PFs (10% in MCDB105 and M199 medium) collected from women before or after ovulation or with a medium (Veh), with or without 100 nM IGF-1R inhibitor (PPP). (B) AIG colony counts of FE25 cells treated with PF collected at different times to the ovulation. Linear regressions of AIG colony number and days post ovulation are showed. *p < 0.05, **p < 0.01

### Ovulatory FF results in the sustained transformation activity of PF independent of IGF signaling

We determined whether the sustained transformation activity of PF results from FF. The durability of AIG activity in FF and a 5%–5% FF/PF mix was tested after incubating at 37°C for various durations. As shown in Figure 2, both 5% FF and the FF/PF mix demonstrated sustained AIG activity after incubation for up to 30 days. Compared with FF, the FF/PF mixture showed less variation in AIG colony counts in multiple repeated experiments, suggesting a more stable transformation activity when FF is mixed into PF after ovulation. Furthermore, we determined whether IGF2/IGF-1R signaling is responsible for durable AIG activity by adding an IGF-1R inhibitor. We found a high dependence of IGF-1R signaling for the early AIG activity of day 0 (D0) FF/PF. However, this dependence was substantially reduced for D7-FF/PF and was absent for D30-FF/PF (Fig. 2C). These findings indicate that the sustained transformation activity of FF/PF is independent of IGF-1R signaling.

**Fig 2.**
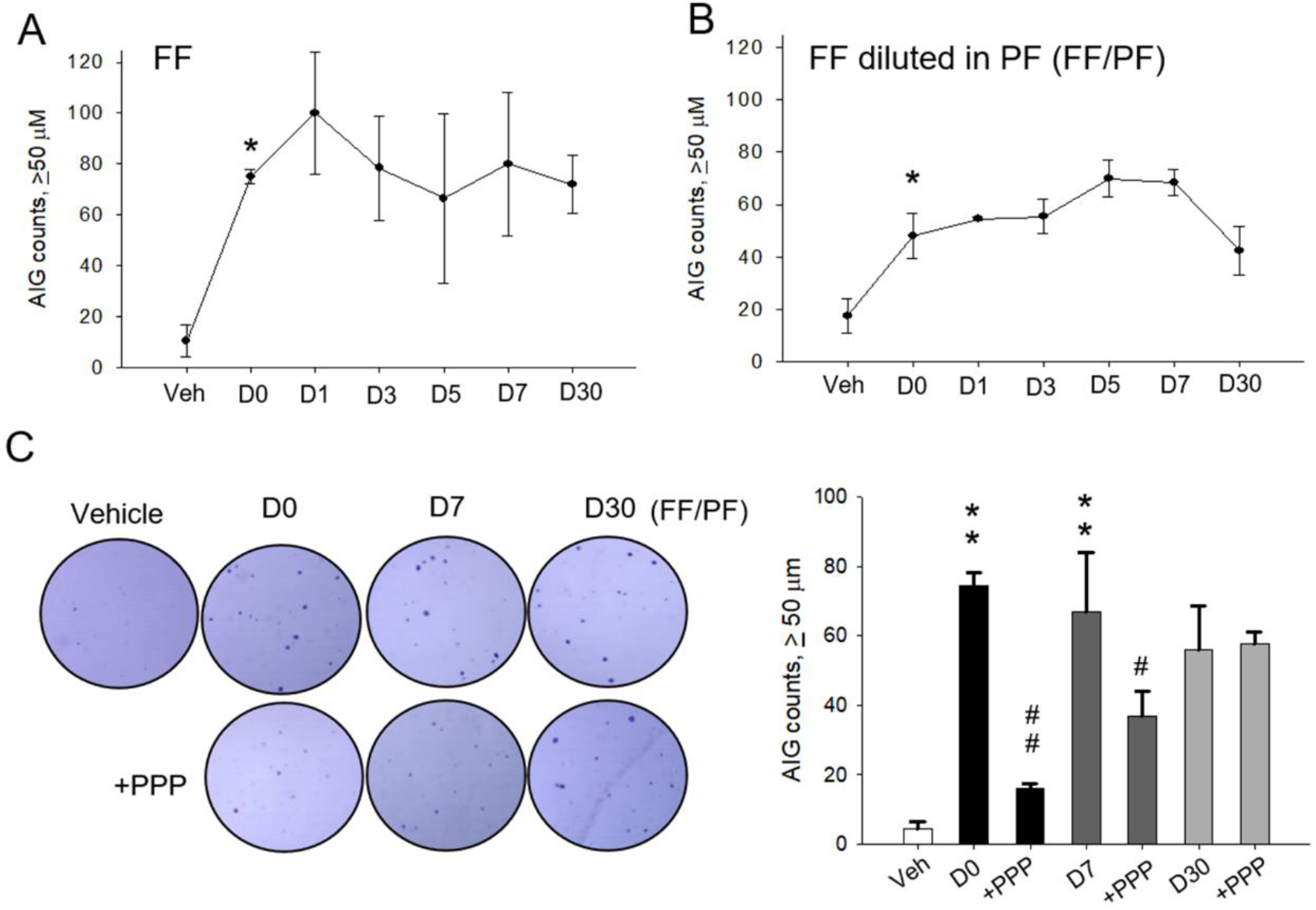
FF sources the sustained AIG activity which is independent of IFG-1R. (A, B) AIG colony counts of FE25 cells treated with a pool of 12 FF samples diluted to 5% with medium (A) or with a 1:1 mix (5% each) of a FF pool and a PF pool (from 15 PF samples shown in Table 1) (B). The fluids were incubated at 37°C for 0 to 30 days (D0, D1, D3, D5, D7, and D30) before being subjected to the AIG assay. (C) Representative AIG images and colony counts of FE25 cells treated with D0-, D7-, or D30-FF/PF with or without 100 nM IGF-1R inhibitor (PPP). * p < 0.05, ** p < 0.01 compared with the vehicle (medium); # p < 0.05, ## p < 0.01 compared with no inhibitor treatment.

### HGF/c-Met transforms FTEC and HGSC in vitro

In our growth assay results of pooled human FF [30852161], we observed a high density of HGF in addition to IGF-axis proteins (Fig. 3A). Expression of HGF was evident in follicular (granulosa) cells surrounding the ovarian follicle (Fig. 3B), and the HGF receptor c-Met was present in the human FTE and HGSC tissue (Fig. 3C). The c-Met protein was also present in FTECs with different extents of transformation (i.e., primarily cultured FTECs, immortalized FTECs with p53 mutation [FT282-V], immortalized FTECs with p53 mutation plus CCNE1 overexpression [FT-CCNE1], and FTECs with p53/Rb disruption [FE25]). The HGSC cell line OVSAHO also expressed c-Met (Fig. 3D). These FTEC cell lines all responded to FF with a marked increase in AIG, which was abolished by inhibiting IGF-1R or c-Met (Fig. 3E). This AIG-promoting activity of FF could be recapitulated to the same level by treatment with IGF2 plus HGF (Fig. 3E). After the same treatments, OVSAHO cells exhibited the same magnitude of the increase in AIG by FF, but the increase was less responded to the treatment of the two receptor inhibitors and was partially recapitulated by pure IGF2 and HGF (Fig. 3E). These results indicate that IGF2 and HGF are responsible for the main transformation activity of FF in FTECs. In fully transformed HGSCs, FF exerts a transformation effect by IGF2, HGF, and additional factors.

**Fig 3.**
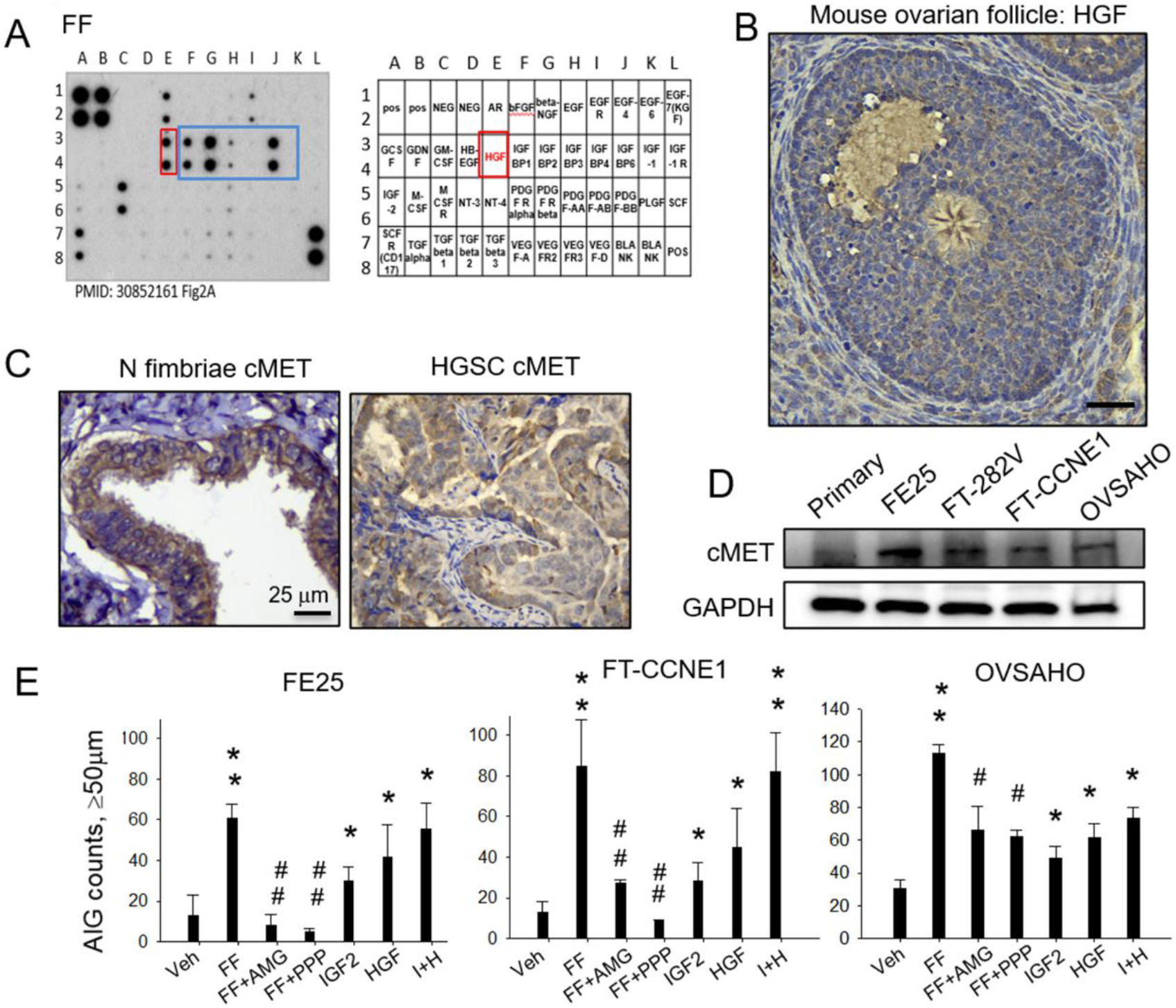
HGF/c-Met, similar to IGF2/IGF-1R, confers the transformation activity of FF in FTEC and HGSC cells. (A) Growth factor array of FF(Hsu et al., 2019) showed abundant HGF (red box) and IGF-axis proteins (blue box). (B, C) IHC of HGF (brown) in the ovarian follicle of a C57BL6 mouse (B), and IHC of c-Met (brown) in the human fallopian tube fimbrium and HGSC tissue (C). Representative images from three independent clinic specimens are presented (scale bar, 25 μm). (D) Western blot analysis of c-Met in different human FTECs and a HGSC cell (OVSAHO). (E) AIG colony counts of FE25, FT-CCNE1, and OVSAHO cells treated with 5% FF, FF plus 10 μM c-Met inhibitor (AMG), FF plus 5 μM IGF-1R inhibitor (PPP), 100 ng/mL recombinant IGF2, 20 ng/mL recombinant HGF, or 100 ng/mL recombinant IGF2 plus 20 ng/mL recombinant HGF (I+H). * p < 0.05, ** p < 0.01 compared with the vehicle (medium with 0.5% DMSO); # p < 0.05 ## p < 0.01 compared with no inhibitor treatment.

### PF-HGF is sourced from FF

We speculated that ovulation sources HGF to PF by draining FF–HGF into the peritoneal cavity. If this is true, then a concentration gradient should be observed from FF and PF to serum. We measured the HGF level in 21 FF and serum samples obtained from women who underwent IVF. Of these samples, 12 were paired FF and serum samples obtained from the same woman at the time of FF/oocyte retrieval. The average concentration of HGF in FF (56 ± 24 ng/mL) was approximately 48 times higher than that in serum (1.16 ± 0.6 ng/mL; Fig. 4A). In the paired samples, a satisfactory correlation (R^2^ = 0.92) was noted between serum- and FF-HGF (Fig. 4B). These results suggest that FF is the primary source of HGF in serum. In addition, in PF collected from the cul-de-sac of 15 women (Table 1), the average HGF level (4.5 ± 4 ng/mL) was approximately 12 times lower than that in FF and approximately 4 times higher than that in serum. Thus, the high concentration of HGF in PF and serum can be attributed to human ovulatory FF.

**Fig. 4.**
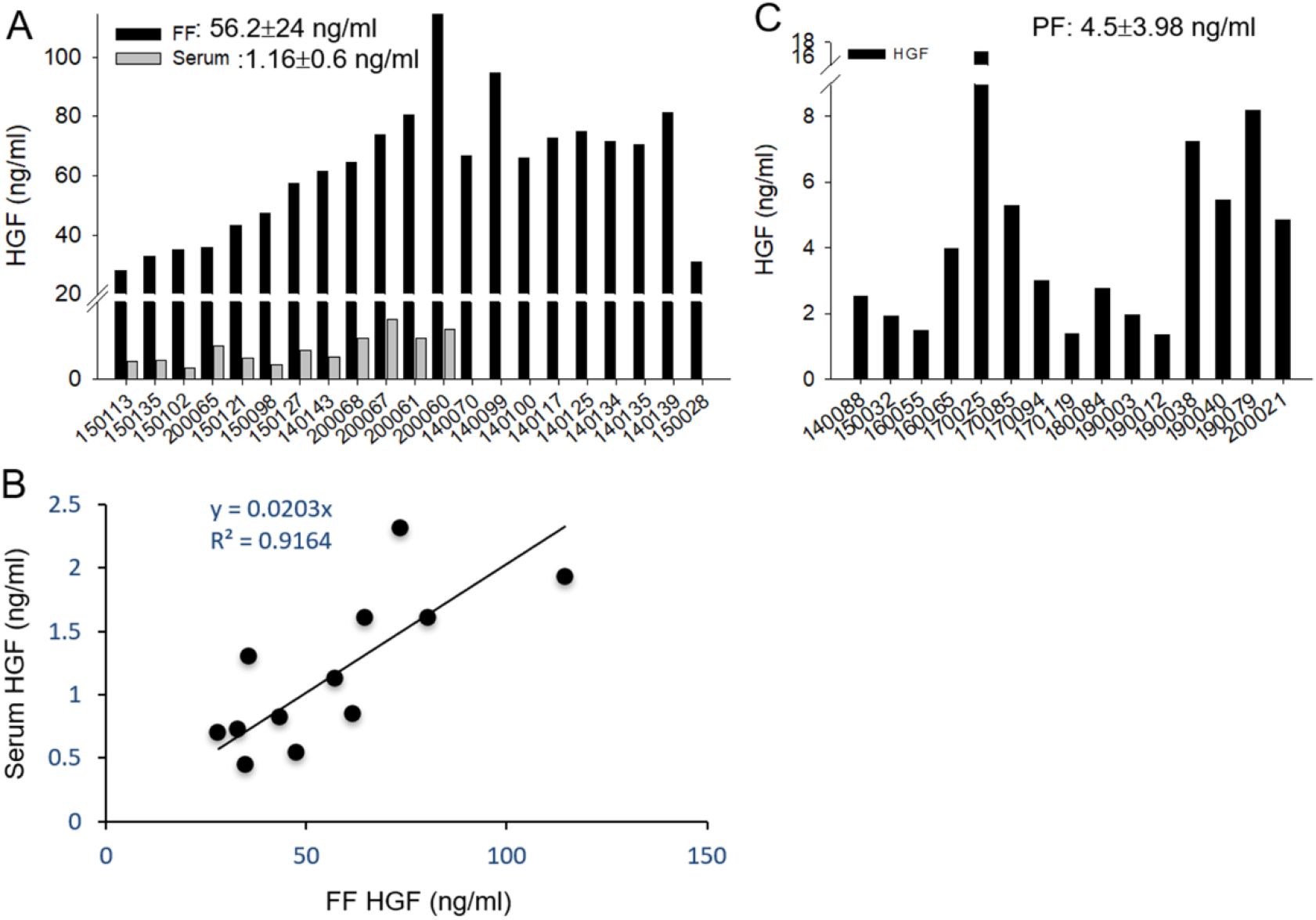
Decreasing concentration gradient of HGF from FF and PF to serum. (A) ELISA analysis of HGF in 21 FF aspirates including 12 paired serum samples collected at the same time. (B) Linear regression analysis of the relationship between the level of FF–HGF and serum HGF in 12 paired samples. (C) ELISA analysis of HGF in 15 PF samples.

### HGF/c-Met signaling is responsible for the sustained transformation activity of PF but does not directly relate to the HGF level

To examine whether HGF/c-Met signaling is responsible for the sustained transformation activity of FF and PF, we conducted the same AIG assay of FE25 cells from PF and FF in the presence or absence of a sublethal dose of a c-Met inhibitor, AMG337 (Fig S1). The treatment readily abolished the AIG activation activity of PF collected either before or after ovulation (Fig. 5A) and the AIG activation activity of the FF/PF after incubation at 37°C for 0, 7, and 30 days (Fig. 5B). In addition, we examined whether the AIG activity of PF correlates with the HGF level in each PF. We observed that the AIG activity of PF was not directly related to the concentration of HGF present within PF (Fig. 5C).

**Fig 5.**
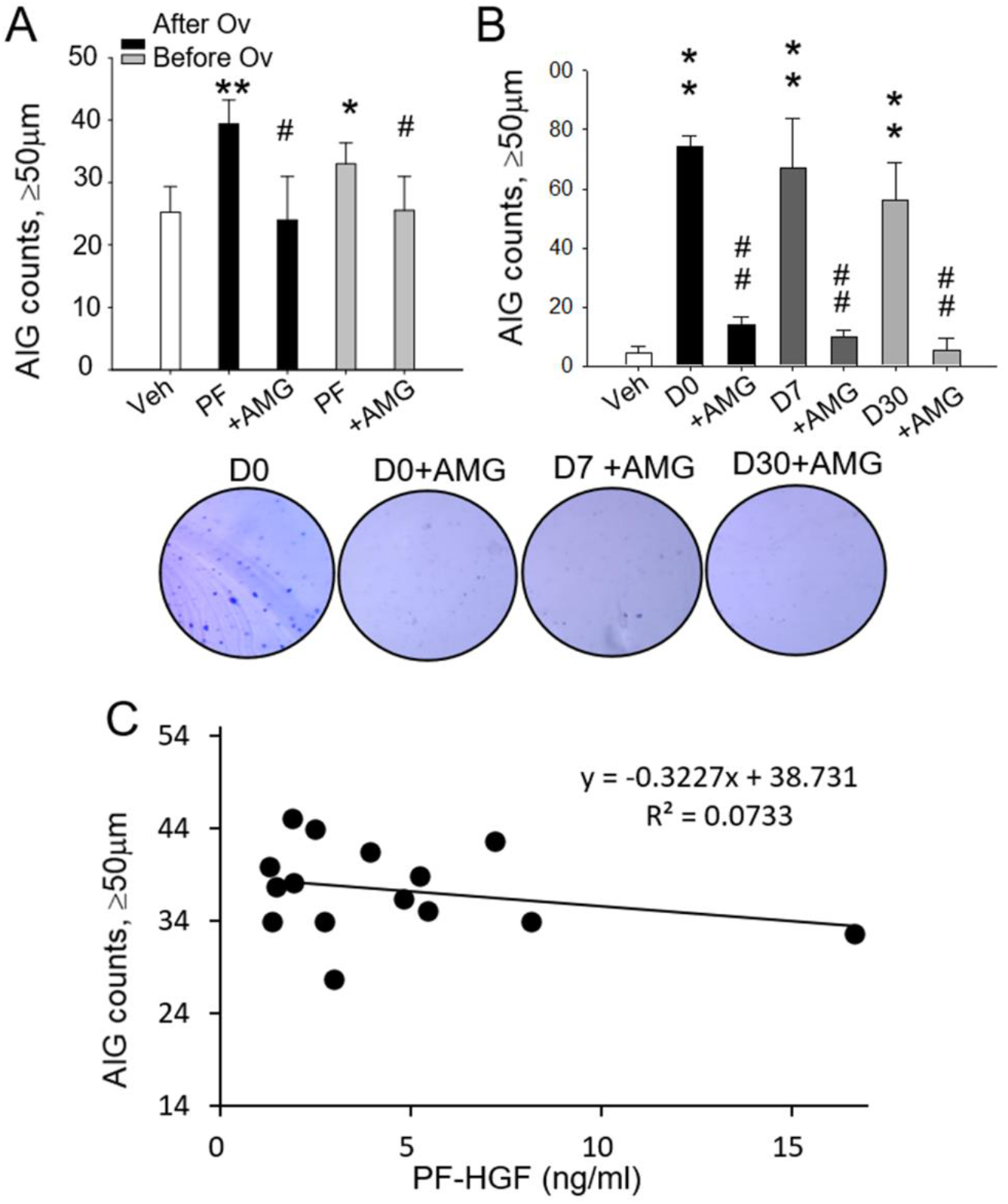
HGF indirectly confers the sustained transformation activity in FF and PF. (A) AIG assay of FE25 cells treated with pooled PF (10%) collected days after (black bar) or before (grey bar) ovulation, with (+AMG) or without 10 μM AMG337, a c-Met inhibitor. (B) AIG assay of FE25 cells treated with FF/PF mix preincubated at 37°C for 0 (D0), 7 (D7), or 30 (D30) days, with or without 10 μM AMG337. Representative colony growths in soft agar are shown. * p < 0.05, ** p < 0.01 compared with the vehicle (medium with 0.5% DMSO); # p < 0.05, ## p < 0.01 compared with FF or PF without inhibitor treatment. (C) Linear regression analysis of the relationship between AIG colony number and HGF level in each PF.

### The coagulation–HGFA–HGF cleavage cascade is responsible for the activation of HGF in FF and PF

HGF is synthesized and secreted as an inactive pro-form that relies on HGFA, a serine proteinase, for cleavage activation. In addition, HGFA requires upstream proteases, such as thrombin, for cleavage and activation (Shimomura et al., 1993). Thus, the activation of HGFA/HGF is linked to the coagulation cascade of proteinases (Fukushima et al., 2018; Miyazawa, 2010; Miyazawa et al., 1996) (Fig. 6A). To examine the presence of this coagulation cascade, we treated the FF/PF pool with tissue factor to activate the extrinsic coagulation pathway. A fibrin clot readily formed after treatment (Fig. 6A). The coagulation factor–depleted supernatant no longer showed the AIG activation activity. Treatment with a thrombin inhibitor, Dabigatran, resulted in a still lower AIG of treated cells (Fig. 6B).

**Fig 6.**
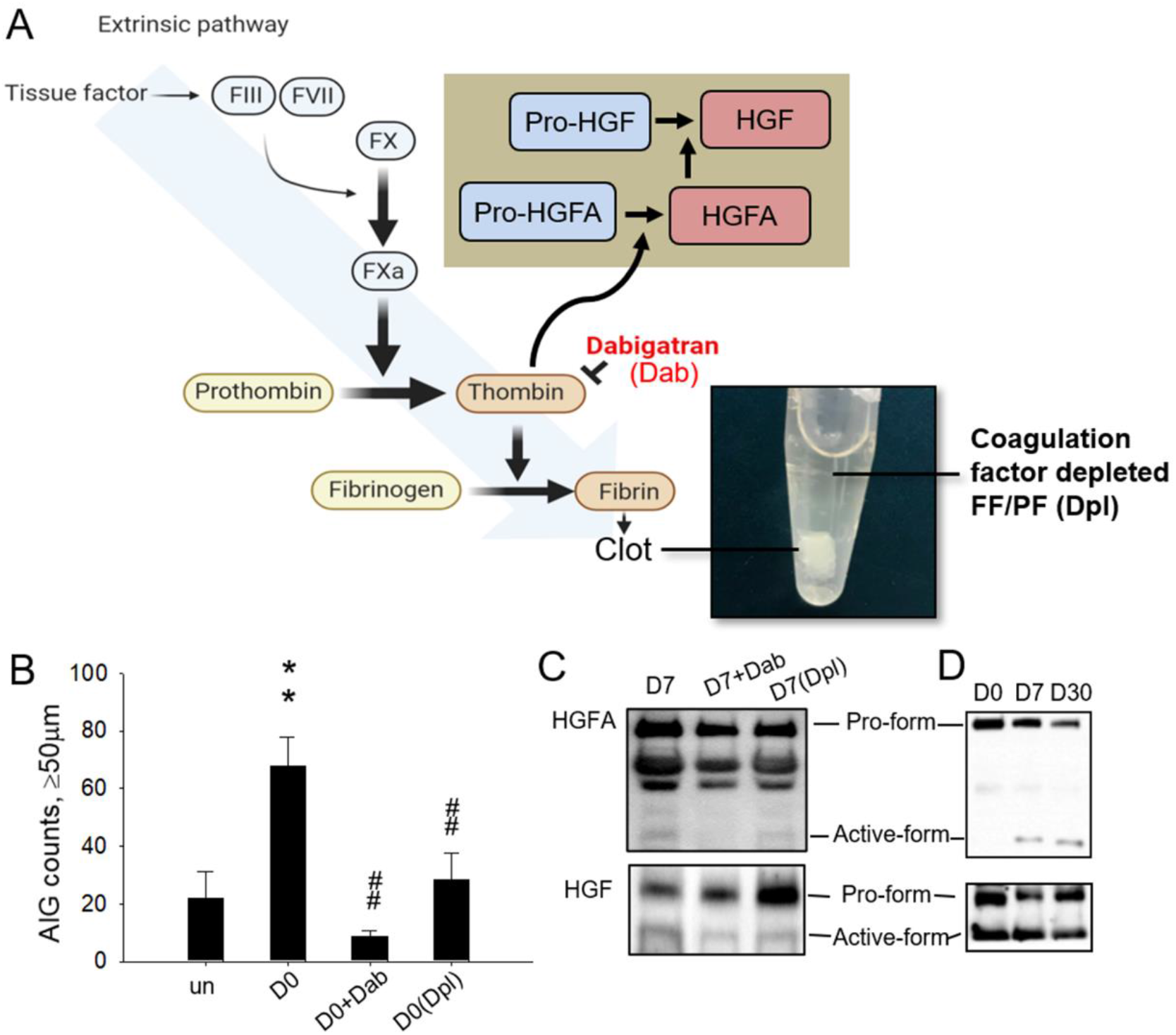
The coagulation–HGFA–HGF cleavage cascade supports the sustained supply of active-form HGFA and HGF in FF/PF. (A) Schematic of the extrinsic pathway linking coagulation to the activation of the HGFA–HGF axis. After initiation by overnight treatment with tissue factor (INNOVIN at 1:1 ratio), a fibrin clot formed in FF/PF, resulting in the coagulation factor–depleted supernatant (Dpl). (B) AIG assay of FE25 cells treated with D0-FF/PF mix with or without a thrombin inhibitor (2 μM Dabigatran) or Dpl. ** p < 0.01 compared with the vehicle (medium with 0.5% DMSO). ## p < 0.01 compared with no inhibitor/depletion treatment. (C) Western blot analysis with pro- and active-form HGFA and HGF in D0-, D7-, and D30-FF/PF and D7-FF/PF with or without 2 μM Dabigatran or with Dpl.

We performed Western blot to examine the cleavage of HGFA and HGF in FF/PF before and after coagulation depletion or thrombin inhibition. As shown in Figure 6C, both treatments resulted in a decrease in active-form HGFA/HGF and an increase in pro-form HGF, indicating a decrease in the coagulation–HGFA–HGF cleavage cascade. In addition, we observed the cleavage of HGFA and HGF pro-forms in FF/PF after incubation at 37°C. A low level of active-form HGFA was observed on day 0, which increased after 7 days of incubation and was maintained on day 30 (Fig. 6D). At the same time, a progressive consumption of pro-form HGFA was noted, and the levels of pro- and active-form HGF were maintained at high levels throughout (Fig. 6D). Thus, an abundant reserve of coagulation–HGFA–HGF cascade proteins support a sustained supply of active-form HGF. After 30 days, the reserve of pro-form precursors remained high, suggesting an even longer supply of active-form HGF.

### The coagulation–HGF cascade of FF/PF plays a significant role in the transformation of FTECs and EOCs in vivo

Given the critical role of the coagulation cascade in HGF activation and cell transformation, we performed xenograft tumorigenesis assays to validate the transformation role of FF/PF and the coagulation cascade. The nontumorigenic FE25 cells were subcutaneously injected into NSG mice, together with the 5% FF/PF with or without coagulation depletion or thrombin inhibition. In accordance with the finding of our previous study, in which FF co-injection resulted in tumor growth in 7 of 11 mice (Hsu *et al.*, 2019), we observed that co-injection of FE25 cells with the FF/PF mix caused tumor growth in 5 of 10 mice. By contrast, co-injection of a thrombin inhibitor with treated FF/PF resulted in tumor growth in only 2 of 12 mice, and co-injection with coagulation factor–depleted FF/PF resulted in tumor growth in only 1 of 6 mice. Moreover, co-injection with recombinant HGF caused tumor growth in 3 of 9 mice (Fig. 7A). Given that not only FTEC but also HGSC cells express c-Met and respond to HGF for AIG, we examined the role of thrombin in ovarian tumorigenesis by using an orthotopic tumorigenesis model in which OVSAHO cells, an HGSC cell line, were injected into the ovarian bursa of NSG mice. The thrombin inhibitor Dabigatran was orally administered on day 0 and twice weekly for 30 days. Significant amelioration of ovarian tumorigenesis was observed after 5 months, with the average tumor weight reduced from 1522 ± 768 to 560 ± 703 mg (p = 0.01; Fig. 7B). Furthermore, in an orthotopic syngeneic transplantation model, a non-HGSC type mouse EOC cell line, ID8(Walton et al., 2016), was orthotopically injected in C57BL6/J mice. The same treatment with a thrombin inhibitor resulted in a significant decrease in tumorigenesis after 4 months, with an average tumor weight of 110 ± 64 mg compared with 460 ± 198 mg (p = 0.002) in the nontreated group (Fig. 7C). The in vivo analysis further supported that the coagulation cascade plays an essential role in the malignant transformation of FTECs and the development of EOC.

**Fig 7.**
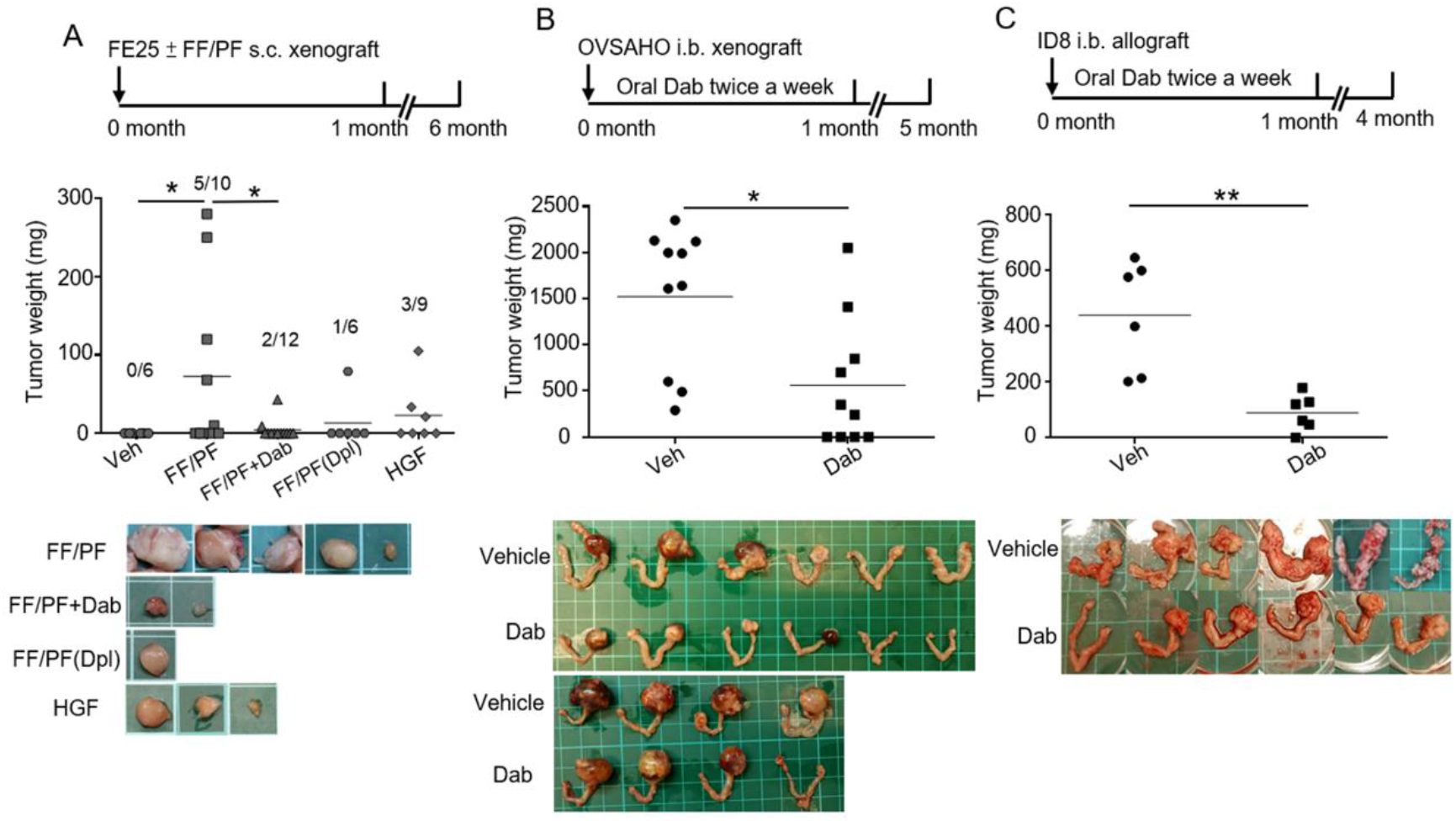
The coagulation–HGF cascade in FF/PF plays a major role in the transformation of FTECs, HGSCs and other EOC cells in vivo. (A) In the xenograft tumorigenesis of FE25 cells in NSG mice, 2 million FE25 cells were subcutaneously injected together with 200 μL of medium and 5% FF/PF with or without pretreatment with the thrombin inhibitor Dabigatran (Dab) or coagulation factor–depleted supernatant (Dpl) or with 20 ng/mL recombinant HGF (100-39H, PeproTech). Tumor growth was observed at 6 months. (B) Orthotopic xenograft tumorigenesis of OVSAHO cells in NSG mice. OVSAHO cells (2 million) were injected into the ovarian bursa (i.b.). Dabigatran was orally administered twice weekly for 1 month, and mice were sacrificed after 5 months. (C) Orthotoic allograft of mouse ID8 (non-HGSC type EOC) cells to C57BL6 mice, with or without the same thrombin inhibitor. Ovarian tumor pictures and tumor weights are shown. ** p < 0.01 vehicle (ddH_2_O with 5% DMSO) compared with dabigatran treatment.

### FF–HGF promotes the growth of cells from the ovarian surface, fallopian tube fimbria, and corpus luteum before and after luteinization

We examined the physiological significance of HGF activity in FF and PF. We proposed that, in the short term, HGF may be responsible for repairing the ruptured ovarian surface and the fimbrial epithelium injured by ovulatory reactive oxygen species (ROS). In addition, after luteinization and ovulation, the ovarian follicle transforms into the corpus luteum. The development of corpus lutein (CL) usually starts with a central hemorrhage (i.e., corpus hemorrhagica), resulting in vigorous coagulation activation. CL has a life span of 7–14 days and plays the physiological role of secreting progesterone to support embryo implantation. We speculated that the high concentration of FF–HGF supports the long-term growth of CL through the same coagulation–HGF cascade. Fig. 8A shows the high expression of c-MET in the CL cells, oviductal epithelium, and OSE. We investigated the effect of FF–HGF on cells derived from these three sites. As shown in Fig. 8B, D0-FF/PF increased the proliferation of KGN granulosa cells with or without luteinization, and the increases were diminished by the inhibition of either IGF-1R or c-MET. However, the D30-FF/PF-induced proliferation could only be abolished by c-MET inhibition (Fig. 8B). Activation of c-MET signaling in KGN cells was evident in Western blot analysis (Fig. 8C), showing an eight-fold increase in c-MET phosphorylation after FF/PF treatment. Treatment with a thrombin inhibitor or coagulation factor–depleted FF/PF reduced phosphorylation by 32% and 71%, respectively. The same mechanism of FF/PF-induced proliferation and c-MET signaling was observed in luteinized KGN cells, with the same magnitude of changes.

**Fig 8.**
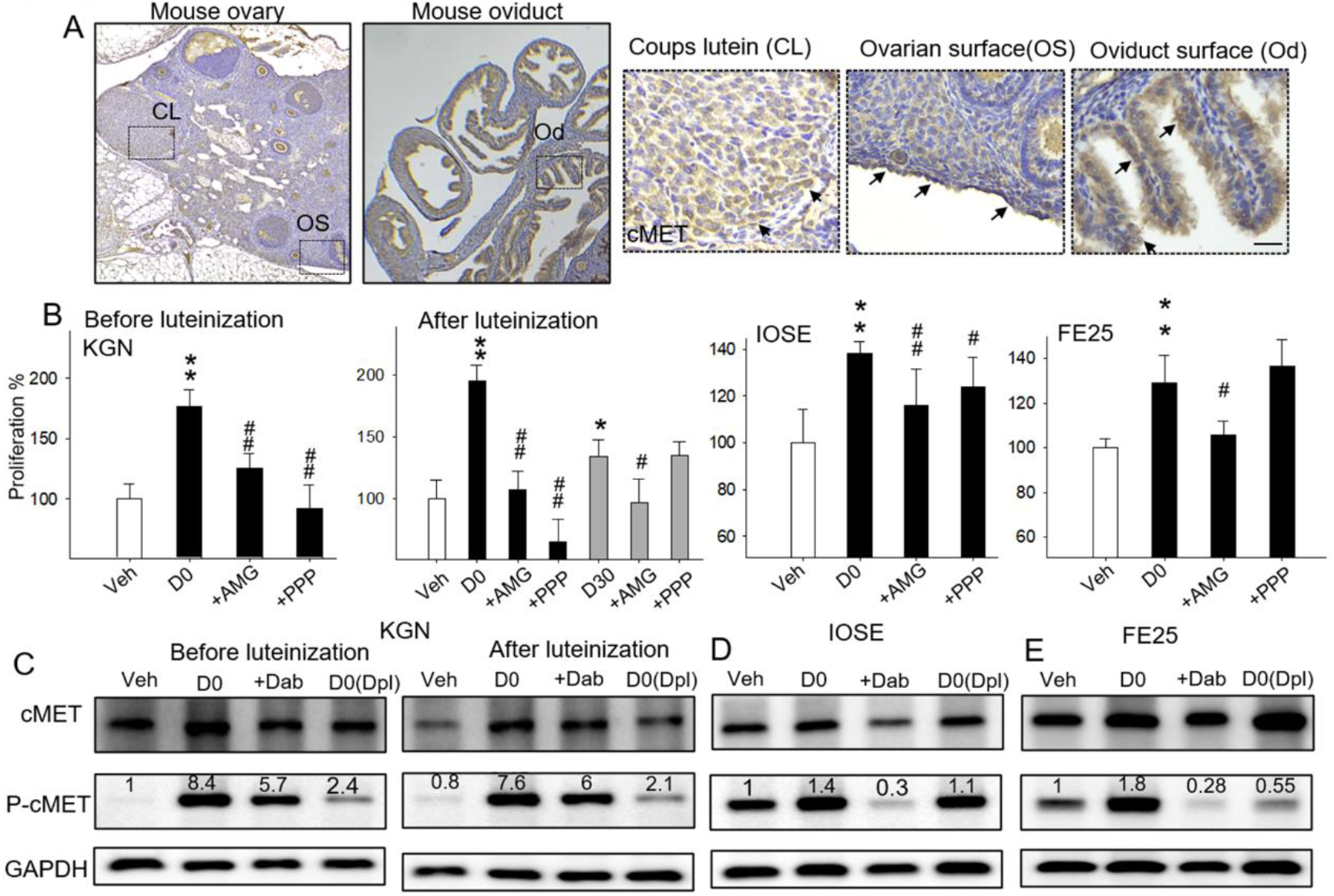
The coagulation–HGF cascade and c-MET signaling supports the proliferation of cells of the corpus luteum, ovarian surface, and fallopian tube epithelium. (A) IHC shows c-Met expression (brown) in the diestrous phase mouse ovary procured 6 h post hCG after PMSG induction. The enlarged picture (dotted box) shows the corpus luteum (CL), ovarian surface, and oviduct surface expression of c-MET (arrows) (scale bar, 25 μm). All images are representative of three independent mice. (B) Proliferation assay of CL cells (KGN) at 72 h with or without 0.6 IU PMSG-induced luteinization as well as ovarian surface cells (IOSE) and fallopian tube fimbrial epithelial cells (FE25), after treatment with 5% D0-FF/PF (D0) with or without 10 μM AMG337 or 100 nM PPP. D30-FF/PF was also tested in luteinized KGN cells. (C) Expression of phosphory- and nonphosphory-forms of c-MET 1h after treatment with FF/PF with or without thrombin inhibition (Dab) or coagulation factor–depleted supernatant (Dpl). * p < 0.05, ** p < 0.01 compared with the vehicle (medium with 0.5% DMSO) of each group. # p < 0.05, ## p < 0.01 compared with no inhibitor used for FF treatment in each group.

Cells derived from the ovarian surface epithelium (IOSE) and fallopian tube epithelium (FE25) also responded to FF/PF with proliferation (Fig. 8D), and the mitogenic effect was diminished by the inhibition of c-MET. Inhibition of IGF-1R also diminished the mitogenic effect in IOSE cells but not in FE25 cells. Unlike KGN cells, both cells exhibited baseline phosphorylation of c-MET, which was moderately increased by D0-FF/PF treatment and markedly decreased by thrombin inhibitor treatment (in both cells) and coagulation factor–depletion (in FE25 cells) (Fig. 8E). The results indicated that the same thrombin–HGF cascade may also be acting in the serum-containing culture of these two cells. Taken together, the results indicate multiple short-term and long-term physiological roles of HGF released through ovulation.

## DISCUSSION

### Identification of HGF as a growth and transforming agent in FF

On the basis of the finding of a previous study that IGF-axis proteins are important transforming and regenerating agents in FF(Hsu *et al.*, 2019), this study demonstrated HGF as the second key growth factor in FF. In contrast to the IGF axis, which exerts a short-term effect on exposed fimbria epithelial cells, the growth and transformation effects of FF–HGF are long-lasting and tightly linked to tissue injury and blood coagulation.

HGF/c-Met signaling plays an essential role in tissue regeneration and a versatile carcinogenic role in many cancers including ovarian cancer(Kwon and Godwin, 2017; Lesko and Majka, 2008; Zhou et al., 2008). Enhanced expression of c-Met has been observed in up to 60% of tumor samples of EOC(Ayhan et al., 2005; Di Renzo et al., 1994; Dimartino and Walz, 1977). A high c-Met expression is associated with a high histologic grade and poor prognosis of EOC(Sawada et al., 2007). However, the oncogenic role of HGF/c-MET in the transformation of the FTE, the primary source of HGSCs, remains unknown. The present study revealed a prolonged transforming activity of HGF in FTECs, HGSCs, and other EOCs, and this activity is sourced from ovulation and sustained in PF.

### Ovulation sources HGF in PF and circulation: Clinical implications in blood testing

Ovulation is a monthly event during which FF and oocytes are released into the peritoneum cavity after the rupture of the Graffian follicle. After ovulation release, FF is readily diffused into the peritoneal cavity and mixed with PF. This intraperitoneal diffusion of ovulatory FF has been well demonstrated in studies on highly concentrated ovarian steroid hormones in PF collected at different times of the menstrual cycle. The concentrations of estradiol (E2) and progesterone (P4) in FF were 50 nM and 100 μM, respectively, and those in serum were approximately 1000-fold lower at pM and nM levels, respectively. Koninckx et al. measured E2 and P4 concentrations in PF. A marked increase in E2 and P4 concentrations are noted after ovulation that rapidly declined to a level equivalent to that in plasma after 7 days(Koninckx et al., 1980). The results of this study showed a rapid equilibration of FF–HGF into circulation after release into the peritoneal cavity. The average concentration of HGF in FF was 12 times higher than that in PF and 48 times higher than that in serum. We found a strong correlation (R^2^ = 0.92) between the HGF concentration in FF and that in serum. This concentration gradient and the close correlation of the HGF concentration in FF and serum indicate that ovulatory FF is a primary source of systemic HGF. HGF is present throughout the body including in the circulation(Blumenschein et al., 2012). High levels of circulating HGF were found in patients with various cancers(Aune et al., 2011). Changes in the serum HGF level reflect therapeutic response, progression, metastasis, and survival in cancers including colon cancer, lung cancer, gastric cancer, and malignant melanoma(Matsumoto et al., 2017). Given the discovery that ovulation is the primary source of serum HGF, one should consider the menstrual status while testing a female patient. Choosing a fixed menstrual cycle day for blood sampling is mandatory. The most feasible time is the day after menstruation. Situations of ovulation inhibition (i.e., the use of OCs and lactation) should also be considered.

### Clinical implication of the sustained transformation activity of FF/PF sourced from ovulation

The results of this study revealed a sustained transformation activity of FF-derived HGF, lasting throughout the menstrual cycle. The AIG activity of PF peaked after ovulation and declined modestly and slowly thereafter. The fact that PF promotes the transformation of FTECs and that the transformation activity results from ovulation warrant the implementation of ovulation inhibition in women with a high risk of ovarian cancer. Because both precancerous FTECs and cancerous HGSCs (OVSAHO) and non-HGSC EOC cells (ID8) respond to this transformation activity, the same practice of ovulation inhibition may be applicable to other EOC types, particularly in unilateral, early stage disease when fertility is spared. Moreover, ovulation inhibition may be vital for other cancers prone to intraperitoneal metastasis including endometrial cancers and GI tract cancers where c-Met is known to be overexpressed(Lee et al., 2018; Li et al., 2015; Yu et al., 2013).

### Ovulation releases the protein pool of the coagulation–HGFA–HGF axis from FF to PF supporting the growth or transformation activity of HGF throughout the ovulation cycle

The study results indicated that HGF was mainly present as a noncleaved latent form in FF and relied on a series of serine proteases, including HGFA and coagulation cascade proteins, for cleavage activation. Notably, we observed an abundant presence of these proteins in FF/PF. At the expanse of upstream latent-form proteases, the active-form HGFA and HGF were maintained. This activation machinery was maintained throughout the 30 days of a menstrual cycle, extending to the next cycle of ovulation. Therefore, a system for long-lasting growth and transformation is reserved in the mature ovarian follicle, which is released and activated through ovulatory injury.

### Reservation and timely activation of two transforming growth signals in FF

The two growth or transformation agents in FF are both conserved before ovulatory release and act precisely on the target (i.e., the injured tissue). The IGF2–IGFBP– PAPP-A axis proteins form a complex in FF, reserving IGF2 activity by binding with IGFBPs. After ovulation, the IGFBP-cleaving enzyme PAPP-A is activated by tethering with the glycoaminoglycan on the membrane of fimbrial epithelial cells, releasing IGF2 to bind with IGF-1R in the vicinity. This leads to the timely repair of FF-exposed tissues through stemness activation and clonal expansion(Hsu *et al.*, 2019). In the case of the coagulation–HGFA–HGF cascade, HGF is precisely activated in the presence of hemorrhagic tissue injury, which initiates the extrinsic pathway of coagulation and activates HGF for cell proliferation and transformation. The two systems in FF likely cooperate to repair the injured tissue after ovulation and inadvertently enhance the transformation of the fimbrial epithelium.

### Physiological role

The HGFA–HGF cleavage axis plays an essential role in the development and regeneration of tissues(Birchmeier and Gherardi, 1998; Nakamura et al., 2011). HGF may mediate mitogenic action on the OSE after ovulation. Early studies examining the action of HGF on OSE cell proliferation have reported conflicting results(Gubbay et al., 2004; Harper et al., 1992). This study showed that the OSE expresses the HGF receptor c-MET, which responds to HGF in FF/PF with autophosphorylation and induces cell proliferation. The study also showed that the fallopian tube fimbrial epithelium, which is exposed to and injured by potent inflammatory and ROS, is regenerated by HGF.

In addition to the short-term repair of ovulation-induced tissue injury, the results of this study indicated a long-term physiological role of the sustained HGF axis signal: to support the growth of luteal cells and maintain the corpus luteum. The corpus luteum is a temporary structure that develops from the ovarian follicle after luteinization and ovulation. The corpus luteum is responsible for the production of progesterone to support the endometrium for the implantation of the embryo(Oliver and Pillarisetty, 2021). The study results revealed that c-Met was expressed in the ovarian follicle and corpus luteum. FF–HGF exerted a proliferation effect on the granulosa cells before and after luteinization. This mitogenic effect was also sustainable and reduced after inhibition of c-Met but not IGF-1R. The results indicate that FF–HGF may facilitate the growth of the corpus luteum before the placenta takes charge to secret hCG to replace the role of progesterone.

In summary, this is the first study to reveal that an ovulation-sourced HGF cascade in the peritoneal cavity supports the long-term growth of the corpus luteum and promotes sustained transformation in EOCs including HGSC. The ovulation-sourced HGF may work not only in the peritoneum but also systemically in the circulation.

## ACKNOWLEDGMENTS

The study was supported by grants of the Ministry of Science and Technology, Taiwan (MOST 108-2314-B-303 -005 -MY2; MOST 107-2314-B-303-013-MY3), and Buddhist Medical Foundation, Taiwan (TCMMP108-01-01; TCMMP108-01-02). The authors acknowledge the core facilities provided by the Advanced Instrumentation Center of Department of Medicine Research, Hualien Tzu Chi Hospital, Buddhist Tzu Chi Medical Foundation, Hualien, Taiwan.

## AUTHOR CONTRIBUTIONS

TYC, HSH, and PCC conceived the project, and TYC, HSH, and SCC obtained funding. TYC, HSH, and PCC designed experiments; HSH, CYH, and MHL generated data. TYC, HSH, MHL, and SCC interpreted data. PCC, SCC, and TYC provided essential patient samples and clinical data; HSH, PCC, and TYC prepared the manuscript. All the authors provided a critical review of the manuscript.

---

**Supplementary Fig 1.**
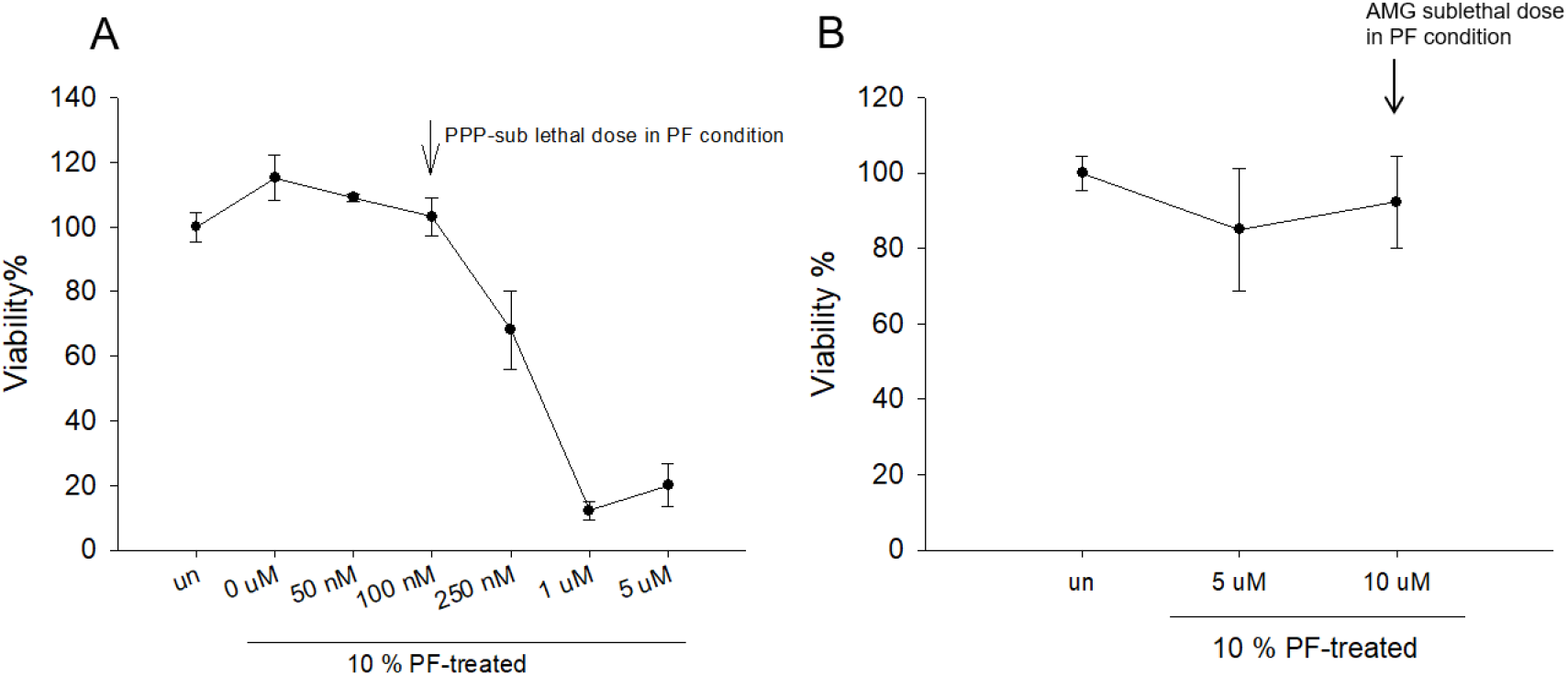
The sub-lethal dose of PPP (A) and AMG337 (B) of FE25 cells under PF treatments

## KEY RESOURCES TABLE

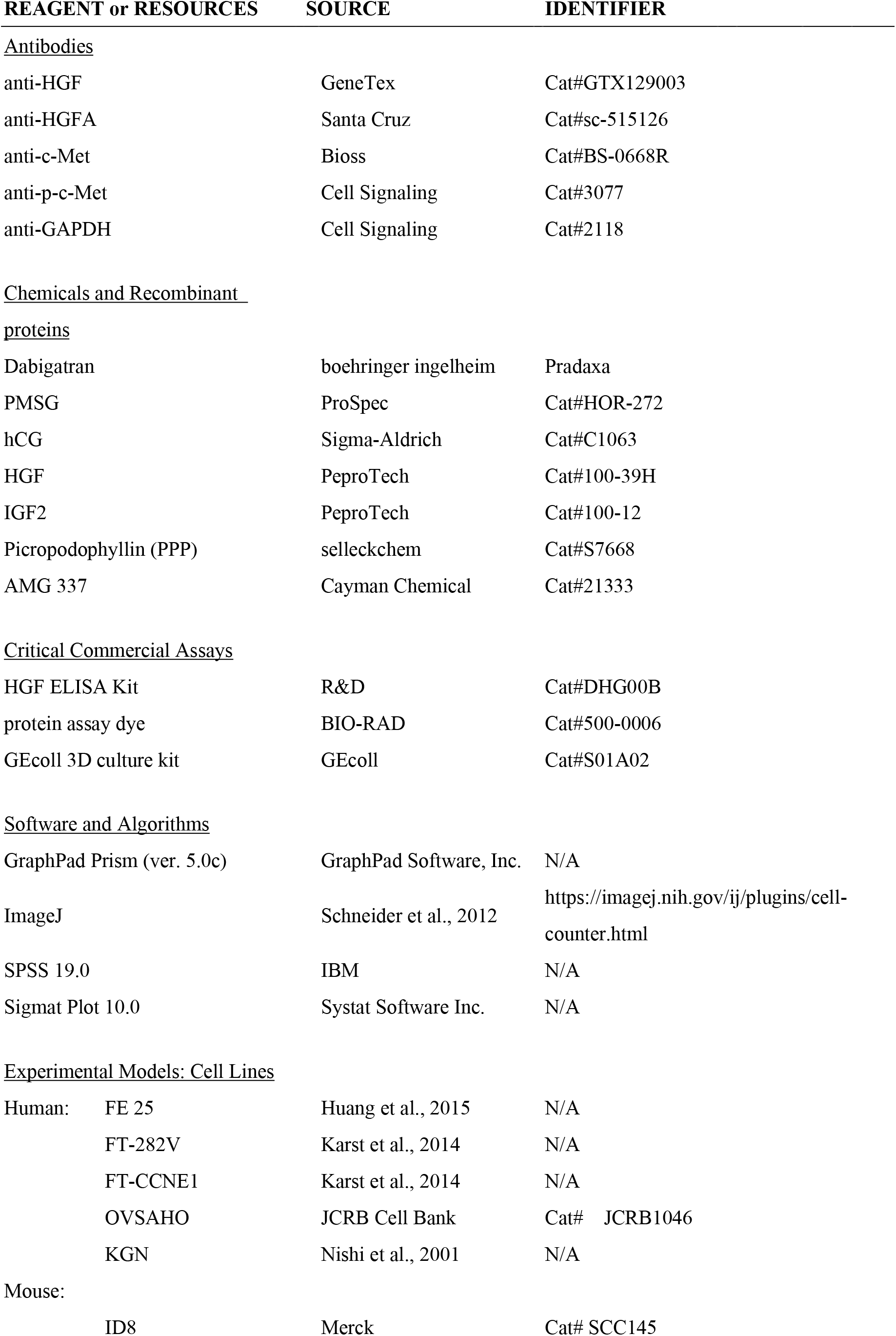
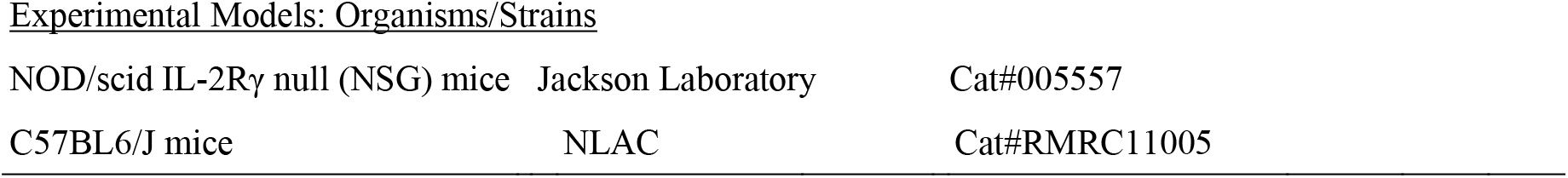

### CONTACT FOR REAGENT AND RESOURCE SHARING

Further information and requests for resources and reagents should be directed to and will be fulfilled by the Lead Contact, Tang-Yuan Chu

## EXPERIMENTAL MODEL AND SUBJECT DETAILS

### METHOD DETAILS

#### Cells

Primary human fallopian tube fimbrial epithelial cells (FTECs) were cultured following the method reported by Paik et al. (Paik et al., 2012) with modifications reported in a previous study(Huang *et al.*, 2015). Three immortalized FTECs were utilized. FE25 cells were established by transducing FTECs with hTERT and HPV16 E6/E7, which inactivates p53/Rb(Huang *et al.*, 2015). FT282-CCNE1 and FT282-V cells were immortalized by hTERT plus a dominant-negative p53R175H mutation, with additional transduction with CCNE1 or empty vector control, respectively(Karst et al., 2014). IOSE is a human ovarian surface epithelial cell line immortalized by using SV40 large T antigen(Maines-Bandiera et al., 1992). All cells were maintained in MCDB105 and M199 media (Sigma) supplemented with 10% fetal bovine serum (FBS) and P/S. The human HGSC cell line OVSAHO was cultured in RPMI-1640 medium with 10% FBS, 100 IU/mL of penicillin, and 100 μg/mL of streptomycin. The mouse epithelial ovarian cancer cell line ID8 was derived from the ovarian surface epithelium and spontaneously transformed through prolonged in vitro culture(Roby et al., 2000). ID8 cells were cultured in high-glucose Dulbecco’s modified Eagle’s medium (DMEM) supplemented with 4% FBS, 5 μg/mL insulin, 5 μg/mL transferrin, 5 ng/mL sodium selenite, 100 IU/mL penicillin, and 100 μg/mL streptomycin. KGN is a nonluteinized human granulosa cell line derived from an ovarian granulosa cell tumor(Nishi et al., 2001). Luteinization was induced by adding 0.6 IU of PMSG. KGN was cultured in DMEM/F12 medium supplemented with 10% FBS, 100 IU/mL of penicillin, and 100 μg/mL of streptomycin.

#### Serum, FF, PF, and tissue specimens

Serum was collected from eight adult women who underwent health examination. Paired serum and FF were collected at the time of oocyte retrieval from 12 women who underwent in vitro fertilization (IVF) as described previously(Hsu *et al.*, 2021; Hsu *et al.*, 2019; Huang *et al.*, 2015; Yeh et al., 2016). To prevent the contamination of the flush medium (which appeared pink due to the presence of phenol red) and blood in FF samples, we discarded the pink and red aspirates. Only yellow FF aspirates were collected. PF was collected from the cul-de-sac of 15 premenopausal women who underwent laparoscopic surgery due to benign, noninflammatory causes. To reduce contamination from blood or tissue damage, PF in the cul-de-sac was aspirated upon entering the peritoneal cavity before initiating surgery. After centrifugation, the supernatants of FF and PF were collected, aliquoted, and frozen before use. The normal fallopian tube tissue was collected from women who underwent opportunistic salpingectomy(Ding et al., 2018), and the ovarian HGSC tissue was collected during debulking surgery. The specimen collection procedures were approved by the Institutional Review Board of Tzu Chi Medical Center, Taiwan (Approval IRB-106-07-A, IRB108-12-A).

#### FF and PF treatment

To simulate the longevity state of FF after drainage into PF after ovulation, we prepared a 5% FF/PF mix in phosphate-buffered saline (PBS) from pooled FF and PF samples. The FF/PF mix was incubated at 37°C, and aliquots were collected after 0, 1, 3, 5, 7, and 30 days, named as D0-, D1-, D3-, D5-, D7-, and D30-FF/PF, respectively. To deplete coagulation factors from D0-FF/PF, a commercialized recombinant tissue factor solution (1×; Innovin, Dade Behring) was added at equal volume. A fibrin clot formed after overnight incubation at 37°C, leaving the supernatant as coagulation factor–depleted FF/PF, named as D0 (Dpl).

#### AIG assay

The AIG assay was performed using the GEcoll 3D culture kit (GEcoll, S01A02). Following the manufacturer’s instruction, we mixed the gel with cells at 37°C and then seeded 500 cells of 20 μL gel into a 24-well plate. After the gel had solidified and the cells were suspended in the interior, 0.5 mL of culture medium was added with or without the tested transforming agent to cover the gel in each well. After two weeks of incubation at 37°C, cell colonies were stained with crystalline violet. Colonies with a size larger than 50 μm were counted.

#### Animal experiments

We used ectopic xenograft, orthotopic xenograft, and orthotopic allograft transplantation mouse models to analyze the transformation activity of FF/PF in vivo. In the ectopic xenograft tumorigenesis model, NOD-*scid* IL-2Rγ^null^ (NSG) mice (Jackson Laboratory) were subcutaneously injected with 2 × 10^6^ FE25 cells in 200 μL of culture medium containing 10% Matrigel plus 5% FF/PF with or without coagulation factor depletion or 5% FF/PF with a thrombin inhibitor (Dabigatran, at a final concentration of 1.26 μg/mL). Mice were sacrificed 6 months later to observe tumor growth. For the orthotopic xenograft or allograft transplantation model, 2 × 10^6^ OVSAHO or ID8 cells, in a 20-μL medium, were injected into the ovarian bursa of NSG or wild-type C57BL6/J mice under a dissecting microscope. For thrombin inhibition, Dabigatran (at a final concentration of 1.26 μg/mL) or a vehicle (0.5% DMSO in ddH2O) was co-injected with cells. Mice were then orally fed with Dabigatran (100 ng/g of body weight) twice a week for 4 weeks. Mice were sacrificed after 4 months in ID8-injected mice and 5 months in OVSAHO-injected mice to observe tumor growth. All experimental procedures were conducted in accordance with the guidelines of the Animal Care and Use Committee of Tzu Chi University (Approval ID: 107-49).

#### Mouse superovulation and ovary sampling

For mouse superovulation, 5 IU PMSG dissolved in 100 μL of PBS was intraperitoneally injected into 8-week-old WT C57BL6/J female mice. After 48 h, 5 IU hCG dissolved in 100 μL of PBS was injected intraperitoneally. At the diestrus phase or 6 h after hCG, the ovaries were surgically removed and subjected to immunohistochemistry (IHC) analysis.

#### Western blot and immunohistochemistry analysis

In Western blot analysis, the protein concentration in the FF or cell lysate was analyzed using a protein assay dye (BIO-RAD, 500-0006). The protein solution was mixed with an equal volume of 2× Laemmli sample buffer and heated at 95°C for 5 min. After cooling on ice, 30 μg of protein was loaded for SDS-PAGE electrophoresis and then transferred onto a nitrocellulose membrane for detection with a specific antibody. Primary antibodies were anti-HGF (GTX129003, GeneTex), anti-HGFA (sc-515126, Santa Cruz), anti-c-Met (BS-0668R, Bioss), anti-p-c-Met (#3077, Cell Signaling), and anti-GAPDH (#2118, Cell Signaling). After washing with Tris-buffered saline with 0.05% Tween, the membrane was treated with appropriate horseradish peroxidase (HRP)–conjugated secondary antibodies and stained with the ECL Western blot detection reagent (GE Healthcare, RPN2209). For the IHC analysis, 4-μm-thick paraffin sections were prepared and stained with the following primary antibodies: anti-c-Met (BS-0668R, Bioss) and anti-HGF (GTX129003, GeneTex). Next, the sections were incubated with appropriate HRP-conjugated secondary antibodies and colorized through 3,3′-diaminobenzidine staining.

## QUANTIFICATION AND STATISTICAL ANALYSIS

Statistical analysis was performed using GraphPad Prism (ver. 5.0c) (GraphPad Software, San Diego, CA, USA), SPSS 19.0, or Excel. Differences between groups were examined using unpaired Student’s t-tests and one-way analysis of variance. The linear correlations of AIG colony numbers with postovulation days and of FF–HGF with serum HGF were measured by calculating the Pearson correlation coefficient.

## DATA AND SOFTWARE AVAILABILITY

This study did not produce additional software. The software used in this paper is publicly available.

